# Real-time observation of replicative helicase assembly onto single-stranded DNA

**DOI:** 10.1101/077800

**Authors:** David Dulin, Zhongbo Yu, Tao Ju Cui, Bojk A. Berghuis, Martin Depken, Nynke H. Dekker

## Abstract

Replicative helicases load onto DNA at the start of replication, and play a vital role by driving the replication fork forward. These helicases assemble into closed multimeric rings that need to encircle single-stranded (ss)DNA to be activated. Though helicase loading on substrates with accessible free ends has been well characterized for the T7 gp4 helicase, a model system for superfamily IV replicative helicases, the physiologically more relevant loading onto exposed ssDNA without free ends remains less well understood. Here, using a label-free assay that exploits changes in the DNA hairpin hopping dynamics to detect gp4 binding and activity, we characterize loading and activation of gp4 on exposed ssDNA without free ends, and find clear evidence of stepwise assembly of the helicase at the fork at physiologically relevant concentrations. The gradual loading onto ssDNA, rather than pre-forming in solution followed by spontaneous ring opening which appears favored at higher concentrations, suggests a new paradigm of stepwise assembly for the helicases in superfamily IV that do not require a separate loading enzyme.

## Introduction

Helicases are essential enzymes that unwind double-stranded DNA (dsDNA) to expose the genetic code for processing by other proteins. These molecular machines power their motion along DNA or RNA through the hydrolysis of nucleotide triphosphates and are classified into six superfamilies according to their structure and direction of motion ^1^. Superfamily IV consists of hexameric helicases involved in bacteriophage and bacterial replication ^2^ (e.g. T7 bacteriophage, T4 bacteriophage, and *E. coli* dnaB), with the gene protein 4 (gp4) of the bacteriophage T7 ^3^ being one of its most studied members. This 63 kDa enzyme is comprised of two domains ^4^: a helicase domain that unwinds the dsDNA template while advancing in the 5’ → 3’ direction, and a primase domain that synthesizes the RNA primers required for lagging strand synthesis ^5^. Structural studies reveal that the helicase domain of gp4 in isolation forms a hexameric ring ^6^, whereas the full length protein adopts a heptameric form ^4^. At high (~µM) gp4 concentrations, the heptameric form is found to be the dominant conformation in the absence of ssDNA, whereas the hexameric form is found to predominate in ssDNA-bound complex ^7^.

During replication of the T7 genome, gp4 is loaded onto ssDNA via a transcription-mediated DNA bubble at the origin. Similar loading platforms are found in T4 bacteriophage, in the ColE1 plasmid of *E. coli*, and in human and mouse mitochondria ^8^. The mechanistic details of gp4 loading have been extensively studied using bulk biochemical approaches, electron microscopy, and X-ray crystallography ^9^. It is known that during initiation no free 5’-DNA end is available to thread through the central channel of a pre-formed gp4 hexamer. Two models have been proposed to lead to the presence of a hexameric form of gp4 encircling the loading site: these invoke the opening of either the hexameric or the heptameric form of gp4 to transfer the ssDNA into the central channel ^7,10^.

Given that the loading of gp4 onto ssDNA is a transient, dynamic, and stochastic event, single-molecule techniques are particularly well suited to its study ^11^. Already, the activity of the T7 replisome has been extensively studied using this type of approach, providing insight into the dynamics of replication over a timescale of minutes with a temporal resolution of seconds ^12^. Higher resolution single-molecule studies (using optical tweezers, magnetic tweezers, and single-molecule FRET) have probed gp4 alone and revealed kinetic details of its DNA unwinding ^13–16^. However, an assay to reveal the dynamics of the underlying loading process has been lacking. Recently, the binding of an antibody onto a chemically modified hairpin stem has been detected by examining changes in the hairpin hopping in an optical tweezers ^17^. Here, using high spatiotemporal-resolution magnetic tweezers ^18^, we utilize an unmodified DNA hairpin stem to observe both the real-time binding kinetics of gp4 on the hopping initial sequence of the stem as well as the subsequent helicase unwinding activity on the remaining part of the stem. This allows us to probe the concentration dependence of the delay between gp4 binding and active unwinding. Quantitative analysis shows that when the available site for loading is shorter than the footprint of gp4, or when gp4 monomers are only present in low and likely close to physiological concentrations, gp4 hexamer forms via gradual assembly onto ssDNA. Conversely, the presence of either a longer ssDNA loading site or higher monomer concentrations facilitate direct loading of a heptamer. These findings thus expose an alternate pathway for the loading of replicative helicases belonging to superfamily IV.

## Materials and Methods

### Experimental assay

The magnetic tweezers apparatus and the hairpin employed here have been previously described ^18^. Briefly, the hairpin handle that is labeled with biotin is attached to a streptavidin-coated MyOne magnetic bead (1 µm diameter, Invitrogen), whereas the other hairpin handle that is labeled with digoxigenin is tethered to the anti-digoxigenin-coated coverslip surface (Fig. 1A). The flow cell is prepared according to the protocol for thin flow cells described in Ref. ^18^.

**Fig. 1:**
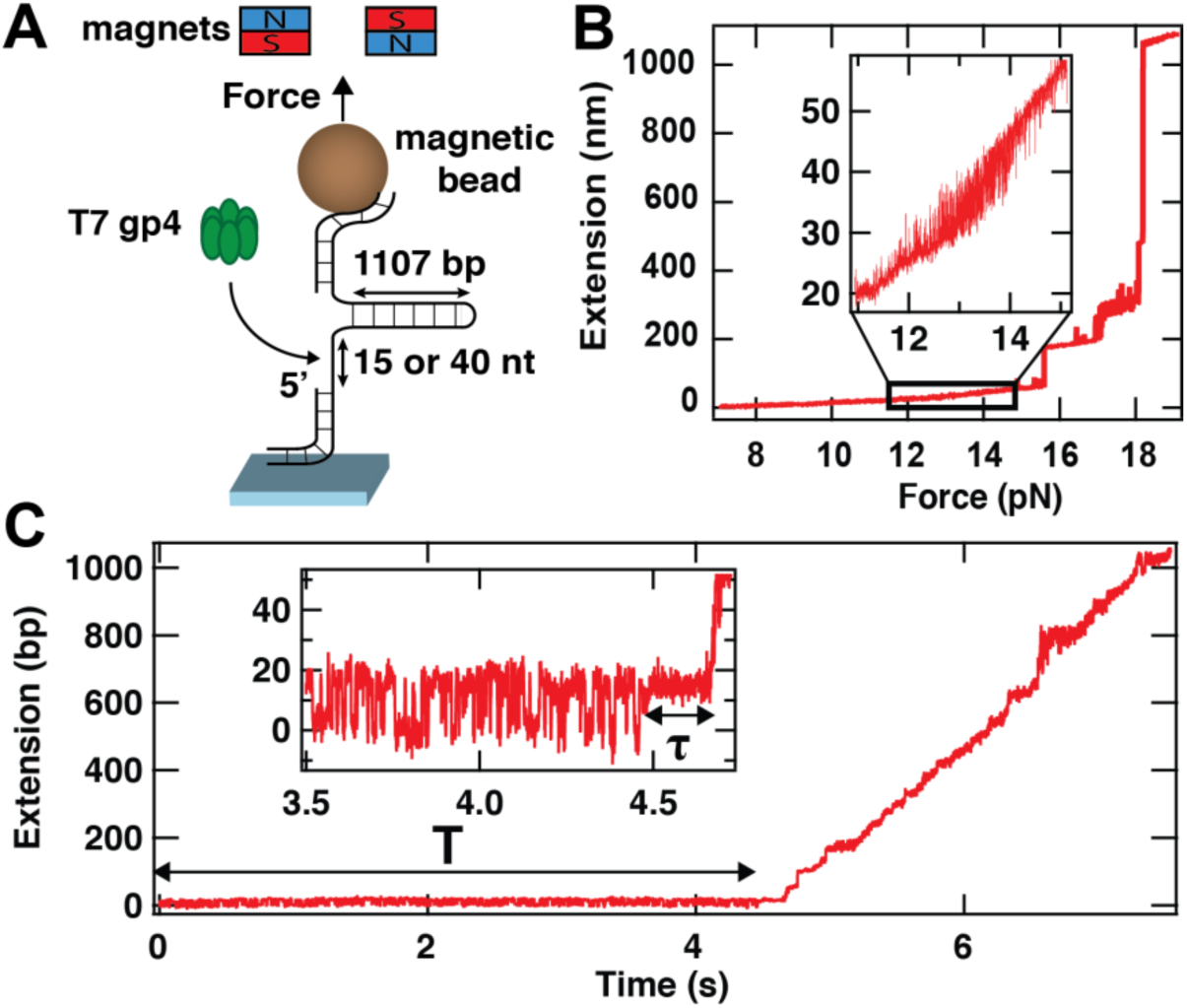
The use of hairpin hopping dynamics as an assay for direct detection of T7 gp4 helicase binding. **(A)** We use a high spatiotemporal-resolution magnetic tweezers apparatus for the observation of gp4 loading and initiation of unwinding. In the instrument, a magnetic bead is tethered to the bottom glass coverslip surface of a flow chamber by a 1107 base pair (bp) DNA hairpin stem. This hairpin has two handles, one of which is partially single-stranded to include a loading site for gp4 of either 15 or 40 nucleotides (nt) in length. Forces are applied to the magnetic bead (and hence to the hairpin) by approaching a pair of magnets to the flow cell surface. **(B)** The extension of the hairpin presented in (A) versus the applied force. The inset shows the force range at which the hairpin hops between a closed state and a state that is partially opened (by ~11 bp)(**Fig. S1B**). **(C)** T7 gp4 loading and unwinding activity on a DNA hairpin stem with a loading site of 15 nt. The inset shows a zoom of the dynamics during ~1 s prior to unwinding. The force applied to the hairpin is set such (~13.5 pN) that the partially opened and closed states of the hairpin are evenly populated (inset) for a time T. At t = ~4.5 s, the hopping is stabilized in the partially opened state for a time τ (inset). Shortly thereafter, unwinding commences and proceeds until the hairpin is fully unwound (~6.8 s).

### Protein concentrations and buffers

We prepare the T7 gp4 helicase (16 µM monomer stock, Biohelix, MA, USA) as detailed in ^13^. Briefly, we first incubate 2 µM of T7 gp4 monomer for 20 min at 22°C in 20 mM Tris-HCl (pH 7.5) buffer containing 2 mM dTTP (Sigma-Aldrich, The Netherlands), 3 mM EDTA, 0.01% Tween 20, and 50 mM NaCl. This helicase solution is diluted to the desired hexameric concentration prior to flushing the reaction solution into the flow chamber. The dilution is performed in a reaction buffer containing 20 mM Tris-HCl pH 7.5, 50 mM NaCl, 7 mM MgCl_2_, 3 mM EDTA, 0.01% Tween-20, and 2 mM dTTP. In the magnetic tweezers experiments, we flush in helicase concentrations of 2, 5 or 10 nM (concentrations are specified in terms of hexameric gp4). This range was delimited by negligible observable activity at lower concentrations (0.75 nM) and a reduction of the start-up time τ to values that were too short to be reliably quantified at higher concentrations (15 nM).

### Characterization of the hairpin and its dynamics in the absence of gp4

Before acquiring data in presence of gp4, we characterize the hairpin and its dynamics as described in **Fig. S1** of the **Supporting Information** and in Ref. ^18^. We verified explicitly that there is no observed correlation between the hopping dynamics and the attachment point of the hairpin to the magnetic bead (**Fig. S1C-D**), as might be expected were the torques generated by off-center attachment to result in hairpin unzipping ^19^. Following this calibration procedure, in the absence of applied force we flush in gp4 at the desired concentration in the reaction buffer described under **Protein concentrations and buffers**. We then ramp up the applied force to the force at which both the closed and partially open states are evenly populated (~ 13.5 pN, Fig. 1B-C). At the end of the acquisition, the force is brought back to 0 pN for ~1 min to completely refold the hairpin and remove any bound gp4. The spatiotemporal resolution of our assay at an acquisition frequency of 500 Hz ^18^ is sufficient to resolve the ~10 nm peak-to-peak steps along the *z*-axis of the hairpin-tethered magnetic bead that result from the hopping of the first ~11 bp at the stem of the 1107 bp hairpin induced by thermal fluctuations (Fig 1B, Fig. 2, 3, 4). The hopping dynamics of this hairpin stem (Fig. 1B) have been previously characterized using identical buffer conditions ^18^ and revealed lifetimes of ~10 ms for both the closed and the open states when both states are evenly populated.

**Fig. 2:**
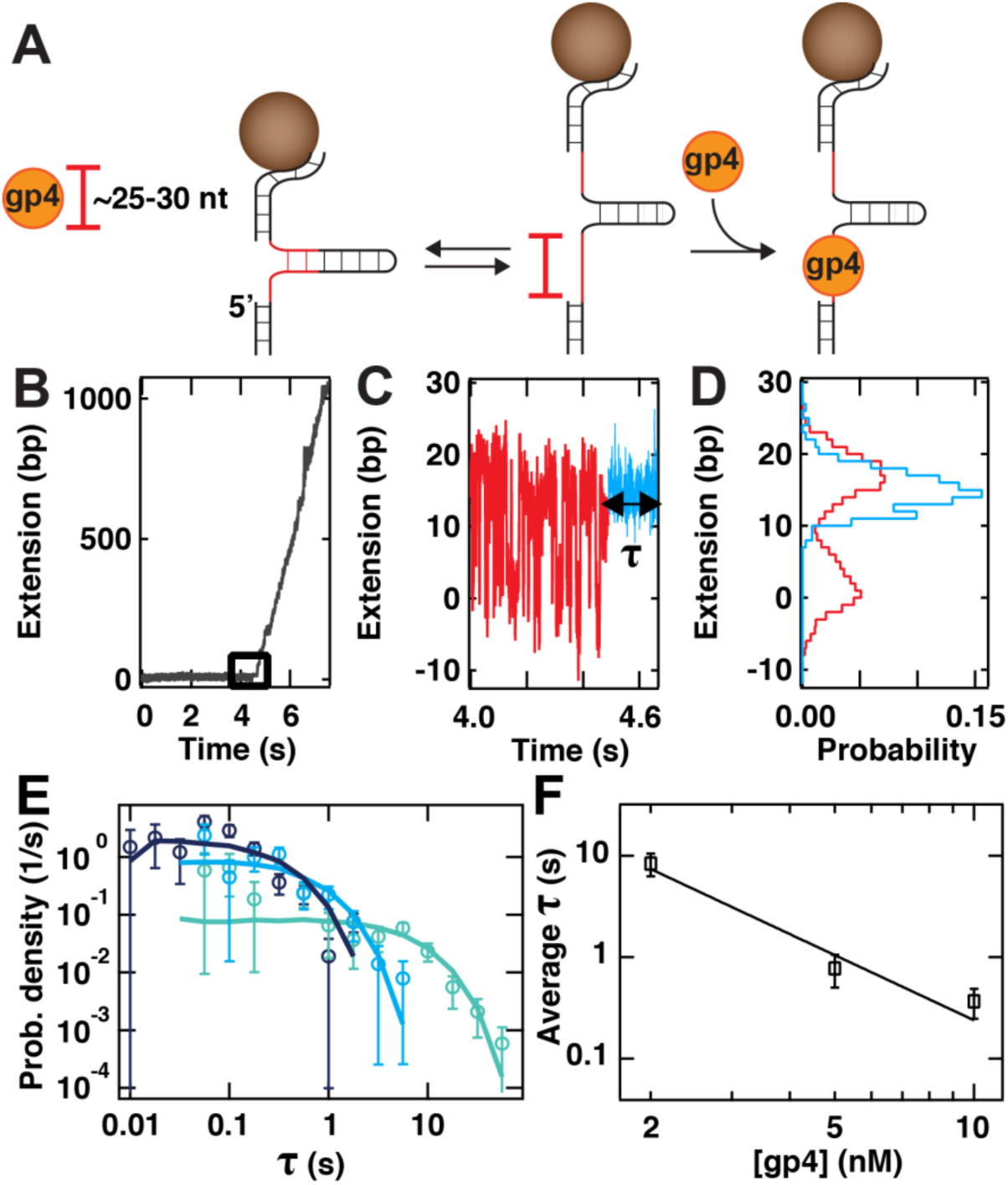
The tight-lock configuration is a signature of a primary binding mode at low gp4 concentrations. **(A)** (Left): The single-stranded loading site adjacent to the DNA hairpin has a length of 15 nt (**Fig. 1A**). As the footprint of gp4 (orange disk) on ssDNA is ~25-30 nt, the protein cannot bind to the loading site when the hairpin is closed. (Center): A small portion (~11 bp) of the hairpin sequence is relatively unstable and unzips under the influence of thermal fluctuations, resulting in transitions between a closed state and a partially opened state (~26 nt total length, red bar) at the applied forces (see inset to Figure 1C). (Right): This provides sufficient space for gp4 to load onto the loading site. In all panels, both the loading site and the unstable portion of the hairpin are indicated in red. **(B-C)** At an applied force of ~13.5 pN, the hairpin hops steadily, evenly populating the closed and partially opened states, until gp4 binds and after some time unwinds the stem of the hairpin. The overall trace is shown in (B), whereas a zoom-in onto the data just prior to helicase unwinding (corresponding to the black square in (B)) is shown in (C). In the zoomed-in trace, the hairpin hopping dynamics (red) are halted and that the partially opened state is stabilized for a time τ (light blue). **(D)** Histograms showing the fraction of time the traces from (C) spent at different positions (same color code as in (C)). The data in panels (B)-(D) were acquired at a gp4 concentration of 5 nM. **(E)** The probability density distribution of the time τ between binding and unwinding for the tight-lock configuration at increasing gp4 concentrations: 2 nM (N = 40, turquoise), 5 nM (N = 63, cyan) and 10 nM (N = 67, dark blue). The error bars represent the one standard deviation confidence intervals obtained from 1000 bootstraps. The solid lines are stochastic simulations for a single exponential process (10000 points) with an exit rate extracted from a MLE fit to the experimental data. **(F)** The mean τ extracted from the MLE fits in (E) as a function of gp4 concentration. The solid line is a fit to 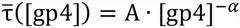 with A = (32.4 ± 14.8) mol^α^ ∙ s and α = 2.1 ± 0.3. The error bars represent one standard deviation confidence intervals extracted from 1000 bootstraps.

**Fig. 3:**
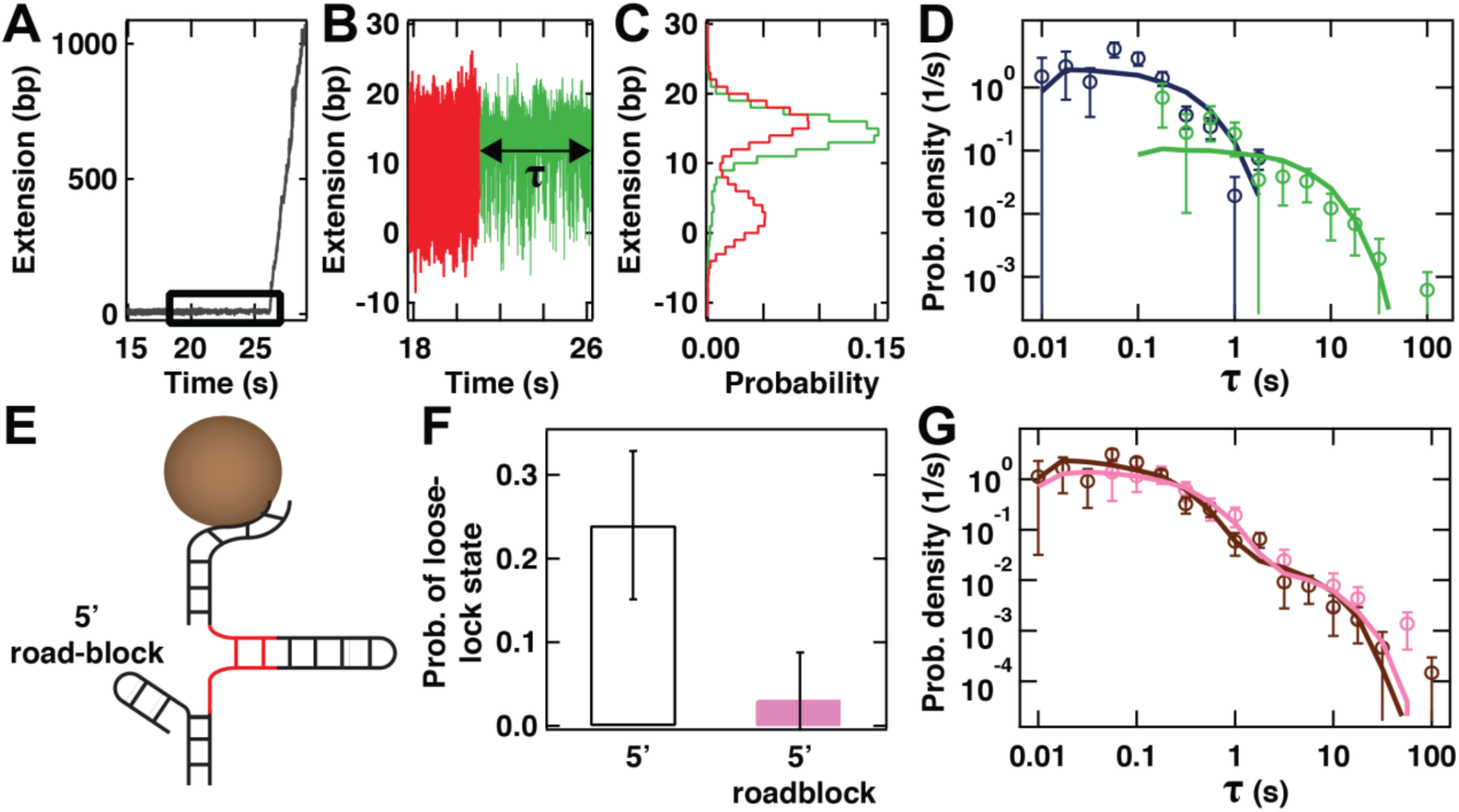
The loose-lock configuration is the signature of a secondary binding mode at higher gp4 concentrations. **(A)** Unwinding of the stem of the hairpin upon binding of gp4 but in a situation **(B)** in which the hairpin hopping (red) is followed by loose locking upon binding (green) for time τ before unwinding. **(C)** Histograms showing the fraction of time the traces from (B) spent at different positions (same color code as in (B)). The closed state of the hairpin is less frequently explored than the partially opened state (red histogram versus green histogram). The data in panels (A)-(C) were acquired at a gp4 concentration of 10 nM. **(D)** The distributions of τ for the tight-lock (dark blue circles, N = 67) and the loose-lock configurations (green circles, N = 21) at 10 nM gp4 concentration and ~13.5 pN. The solid lines are stochastic simulations for a single exponential process (10000 points) with an exit rate extracted from MLE fits to the data. The error bars are one standard deviation confidence interval from 1000 bootstraps. **(E)** Schematic representation of the hairpin in which the 5’-end of the DNA handle preceding the loading site is terminated by a duplex DNA that forms a road block to prevent backwards sliding of gp4. **(F)** Probability of observing a loose-lock configuration with a hairpin with (pink) and without (white with dark blue frame) a dsDNA duplex 5’-roadblock as described in (E) for a gp4 concentration of 10 nM. The error bars represent 95% confidence intervals. **(G)** The full distribution of start-up times τ obtained on the hairpin containing a 5’-roadblock formed from dsDNA as described in panel (E) (pink circles, N = 33). This dataset is to be compared with that which results (brown circles, N = 88) from re-merging the distribution of start-up times τ for the tight-lock (blue circles, panel (D)) and loose-lock states (green circles in panel (D)) obtained on an identical hairpin in the absence of the roadblock. The solid lines represent the results of stochastic simulations for a double exponential process (10000 points) with exit rates extracted from MLE fits to the data. The error bars represent one standard deviation confidence intervals obtained from 1000 bootstraps.

**Fig. 4:**
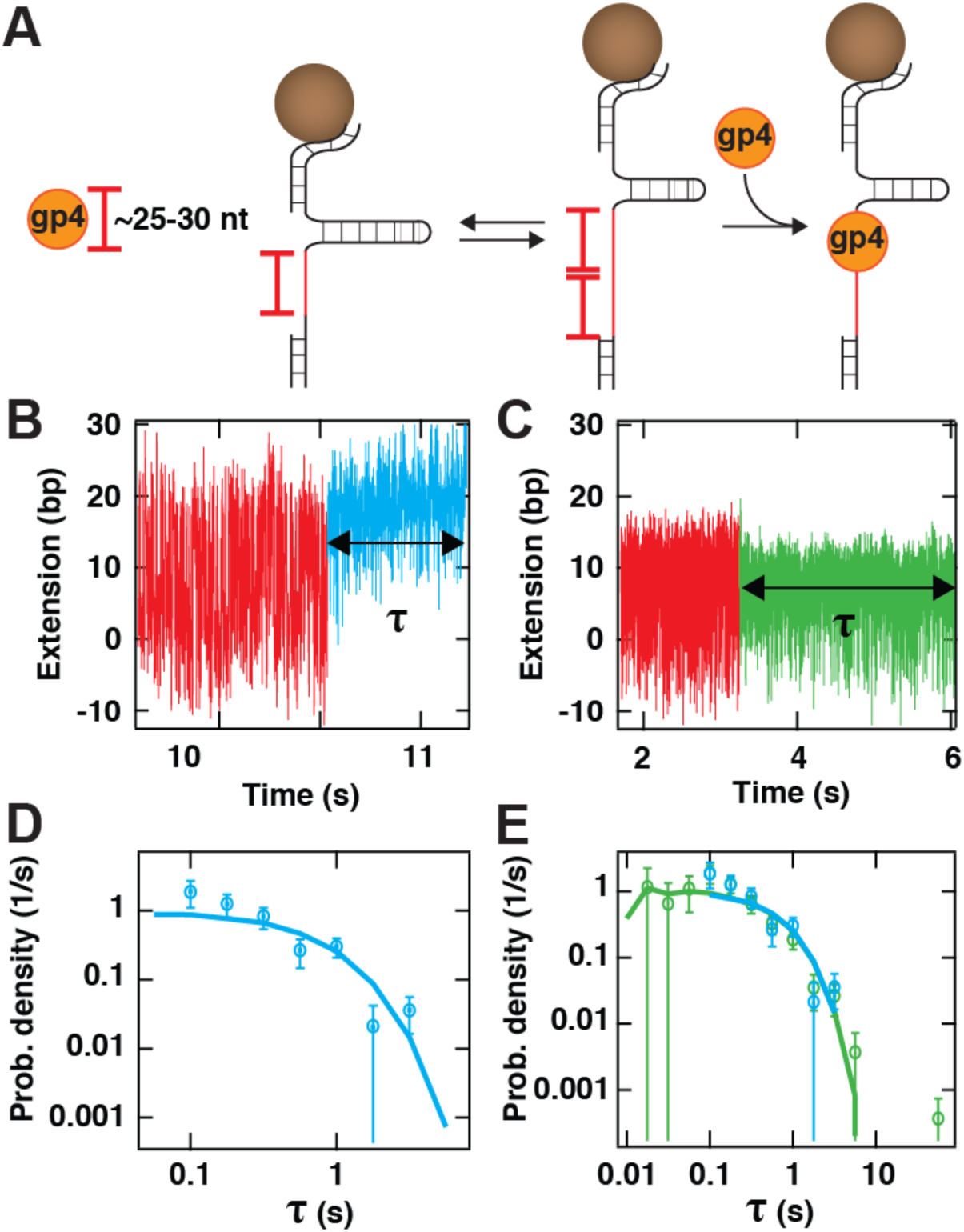
The tight and loose lock configurations of gp4 are conserved on a hairpin with a longer loading site. **(A)** Schematic of the experiment in which gp4 binds onto a 40 nt ssDNA loading site while the hairpin undergoes opening and closing of the first 10 bp of its stem. (Left): When the hairpin hopping dynamics explore the closed state, the loading site offers sufficient space to load a single gp4 (red bar). (Right): When the hairpin hopping dynamics explore the open state, this longer loading site offers sufficient space (~50 nt) to load two helicases. **(B,C)** Following the addition of gp4, we observe that the hairpin hopping dynamics (red) is either (tightly locked in the open state (blue data in panel (B)) or loosely locked (green data in panel (C)) for a start-up time τ prior to unwinding. **(D)** Distribution of the start-up times τ for the tight-lock configuration with a 40 nt ssDNA loading site (blue circles, N = 28). **(E)** Comparison of the distributions of τ for the tight-lock configuration (blue circles, N = 28) and the loose-lock configuration (green circles, N = 33). In panels (D),(E), the solid lines represent a 10000 point simulation for an exponentially distributed stochastic process with a lifetime extracted from MLE fits. The error bars represent one standard deviation confidence intervals derived from 1000 bootstraps. Data in all panels was acquired at a gp4 concentration of 5 nM under an applied force of ~13.5 pN.

### Conversion of the hairpin tether length from nm to base pair

The complete unzipping of the stem of the DNA hairpin results in a change in the end-to-end extension of ~1 µm. To convert this change in extension from a distance unit to a number of unwound bp unit, we determine the bp/nm conversion factor by dividing the full length of the stem of the hairpin (1107 bp, Fig. 1A) by the maximum extension difference (in nm) between the closed hairpin and the fully open hairpin at a given force, as described in Ref. ^14^.

### Determination of the start-up time τ

As exemplified in Fig. 1C, prior to hairpin unwinding by gp4, we observe that the hairpin hopping dynamics is modified for a dwell time τ (Fig. 1C, inset) prior to hairpin unwinding. We determine τ using a thresholding algorithm that was previously used to determine the hairpin hopping dynamics ^18^.

### Identification of tight- and loose-lock binding modes

As described in the main text, the traces acquired in the presence of gp4 are divided into three regimes: prebinding (during which the hairpin hopping dynamics of the hairpin remain unchanged for a time *T*), binding for a start-up time τ, and unwinding. We categorize the mode of gp4 binding as the loose-lock mode if, during the start-up time τ, a double Gaussian function provides a better fit to the data than a single Gaussian function and the probability of visiting the closed state exceeds 0.01. Otherwise, the mode of gp4 binding is identified as the tight-lock binding mode. Representative traces and analysis of the dynamics for the different loading site sizes and gp4 concentrations are shown in **Fig. S3** (tight-lock binding mode) and **Fig. S4** (loose-lock binding mode).

### Characterization of the distributions of start-up times τ

The analysis of the start-up time distribution is explained in more detail in ^20–22^. Shortly, the distribution of τ are described by a probability distribution function with *m* exponentials:

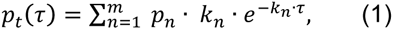

where *k*_*n*_ and *p*_*n*_ are the characteristic rate of the *m*^*th*^ exponential and its probability, respectively. We calculate the maximum likelihood estimate of the parameters (MLE) ^23^ by maximizing:

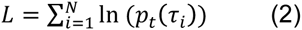

over the parameter set. Here the τ_*i*_ are the experimentally measured start-up times and *N* is the number of collected start-up times τ_*i*_.

## Results

We employ a high-resolution magnetic tweezers assay that exploits magnetic beads tethered to DNA hairpins to monitor protein binding. The hairpins consist of an 1107 bp stem preceded by a ssDNA loading site for gp4 of either 15 or 40 nt in length (Fig. 1A). The sizes of the ssDNA loading site are selected to respectively be smaller and larger than the footprint of gp4 (25-30 nt ^24^). We use the characteristic jump in extension that results from cooperative unwinding of the DNA hairpin at forces exceeding ~16 pN to verify the hairpin nature of the tether (Fig. 1B; **Fig. S1A**). Upon approaching the critical unwinding force, rapid hopping dynamics at the base of the hairpin stem are observed (Fig. 1B), in accord with previous measurements of DNA hairpins using optical and magnetic tweezers ^18,25–29^. These measurements indicate that under the influence of thermal fluctuations, the base of the hairpin stem reversibly opens by ~11 bp. In our experiments, we tune the value of the applied force so that these two states, designated ‘closed’ and ‘partially open’, are equally populated.

Using a hairpin with a 15 nt ssDNA loading site, we observe that data traces progress through three parts upon the addition of gp4: first, the hopping dynamics of the hairpin is unchanged for a time *T* (Fig. 1C); then the hopping dynamics are reduced, with the hairpin appearing locked into a partially open state for a start-up time τ (inset to Fig. 1C); lastly, the hairpin is unwound by gp4 (Fig. 1C). We find that the median time *T* before observing any change in the hopping characteristics of the hairpin (Fig. 1C) depends on the gp4 concentration (**Fig. S2J**): *T*=124 ± 20 s (median ± 95% confidence interval) at 10 nM gp4, and *T*=614 ± 65 s at 5 nM gp4 (measurements of *T* at 2 nM are hampered by constraints on the final size of the data set at high acquisition frequency that limits the duration of experimental recordings 18.) The observed changes in hairpin hopping dynamics are dependent on the presence of gp4, and are, irrespective of the gp4 concentration employed, always followed by the strand separation that is indicative of helicase activity. Therefore, we associate the initial change in hairpin hopping dynamics with the binding of gp4 onto exposed ssDNA for a duration τ (inset, Fig. 1C). Given the restricted size (15 nt) of the loading site presented, gp4 binding is only possible when the hairpin is thermally kicked into its partially open state, which exposes an additional ~11 nt ^18^ to yield a stretch of ssDNA comparable to the gp4 footprint on this substrate (Fig. 2A) ^24^. Binding of gp4 then occupies the available space for the partially opened hairpin, leaving no further room for e.g. helicase sliding and accompanying changes in hairpin dynamics. Consistent with this, we observe that the hairpin hopping dynamics do not change any further (Fig. 2B-D), suggesting that no additional helicases beyond the first one are capable of binding. A histogram of the hairpin extension (Fig. 2D) also shows that there is no significant shift in its maximal extension over the time interval τ, indicating that no unwinding activity has taken place. We coin this binding signature the *tight-lock* configuration (**Materials and Methods**; additional sample traces shown in **Fig. S3**).

### The tight-lock configuration shows the signatures of step-wise assembly of gp4 onto a 15 nt loading site

To further probe the origin of the observed delay between gp4 loading and subsequent unwinding, we probe the start-up time τ at different gp4 concentrations. Distributions of τ for the tight-lock configuration appear exponentially distributed (Fig. 2E) and are well described by single exponential functions (solid lines in Fig. 2E). This yields lifetimes of (8.3 ± *2*.1) s (mean ± standard deviation), (0.8 ± 0.3) s, and (0.4 ± 0.1) s for the 2 nM, 5 nM, and 10 nM helicase concentrations, respectively (Fig. 2F). As only a single gp4 protein has room to bind the loading site prior to unwinding, the decrease in the start-up time τ with increased gp4 concentrations suggests that gp4 does not load onto the ssDNA loading site directly as a complete hexamer. The direct loading of an unwinding-compatible hexamer would predict a concentration dependence only of the time *T* before binding, not in the delay τ between binding and unwinding, in contrast to our observations. This suggests that the initial entry into the tight-lock configuration must be followed the binding of additional gp4 oligomers (of a form to be determined) for the formation of an unwinding-competent gp4. Fitting the concentration-dependence of the start-up times τ to a power law reveals a quadratic dependence (solid line in Fig. 2F), suggesting cooperative binding of gp4 after lock formation. The fact that the start-up times τ are distributed according to single exponential functions (solid lines in Fig. 2E) further suggests a single step to helicase activity after lock formation. Taken together, our data argues for a multi-step binding process, where, after the initial binding, unwinding only happens after a last cooperative binding step completes the hexameric form. Within this model of step-wise gp4 assembly onto ssDNA, the tight-lock configuration also suggests that even an incomplete gp4 oligomer ring binds sufficiently tightly to lock the hairpin into its partially open state.

### A loose-lock configuration suggests binding by a heptameric form of gp4

When examining the 15 nt ssDNA loading site hairpin at the highest gp4 concentration (10 nM), we observe an alternative signature of gp4 binding prior to DNA unwinding **(Fig. 3A-B, Materials and Methods**). Here, too, gp4 binding stabilizes the partially open state (Fig. 3C), suggestive of the presence of gp4 on the loading site. However, we observe that the hairpin appears to continue to explore its closed state, albeit at a lower frequency than during the hairpin hopping dynamics in the absence of gp4 (Fig. 3C). The fact that the hairpin is able to close despite the presence of loaded gp4 implies that the protein must partially overlap the dsDNA handle directly upstream of the loading site. We term this second binding signature the *loose-lock* configuration, as the system predominantly appears to lock into a partially open hairpin state, but occasionally allows the hairpin to close temporarily. We deem helicase dissociation during such hairpin closure to be unlikely, as the time to reenter the locked configuration is much shorter than the original binding time *T*. Additional examples of this binding signature are displayed in **Fig. S4**. In Fig. 3D, we directly compare the distributions of the start-up time τ for the two locked states, and find that the average start-up time τ in the loose-lock binding mode (5.1 ± 2.8) s exceeds that of the tight-lock binding mode (0.4 ± 0.1) s by more than an order of magnitude. This large difference in the start-up times, together with the different binding signatures in the DNA hairpin hopping dynamics, suggests that different helicase configurations are responsible for the loose- and tight-lock binding modes.

Structural studies have shown that at high gp4 concentrations, the formation of a heptameric form of gp4 is favored over the hexameric form. This heptameric form adopts a conformation that binds only poorly to ssDNA, but is sufficiently wide to encircle dsDNA ^7^. We hypothesize that the loose-lock binding mode observed at high gp4 concentrations originates from the increased occurrence of large but poorly binding heptameric gp4, whereas the tight-lock binding originates from the stepwise assembly of hexameric gp4. To test predictions made by this hypothesis, we assessed whether the loose-lock binding mode is compatible with transient sliding of gp4 onto the dsDNA directly upstream of the ssDNA-loading site. To this end, we configured a roadblock at this location (5’ roadblock formed by a short DNA hairpin, Fig. 3E). This roadblock should be sufficiently large to prevent any helicase bound to the ssDNA loading site from sliding backwards ^7^. Under these conditions and 10 nM gp4, we find that the probability of observing the signature of the loose-lock binding mode on the hairpin hopping dynamics is drastically reduced, from (24 ± 9) % (mean ± 95% confidence interval) in the absence of the roadblock (Fig. 2E, N = 88) to (3 ± 5) % in the presence of the roadblock (Fig. 3F, N = 34). The inner diameter of heptameric gp4 is sufficiently large to encircle one, but not two dsDNAs ^7^. Therefore, the observed reduction in the probability of hairpin closure in the presence of gp4 when the short loading site is abutted by a 5’-roadblock supports the identification of the loose-lock binding mode with the loading of the heptameric form of gp4.

To gain further insight into how a heptameric form of gp4 might bind to the loading site, we examined the distribution of start-up times τ for the hairpin including the 5’-roadblock. This distribution (10 nM gp4, pink circles, Fig. 3G) is described by a bi-exponential distribution (pink solid lines, Fig. 3G), which contains a short-lived majority population (probability *p*_1_ = 0.78 ± 0.12 with short *t*_1_ = 0.41 ± 0.12 s) and a long-lived minority population (probability *p*_2_ = 0.22 ± 0.12, *t*_2_ = 9.3 ± 3.5 s). Notably, this distribution overlaps with the result of re-merging (brown circles, Fig. 3G) the distributions associated with the tight- and loose-lock binding modes (shown in distinct colors in Fig. 3D) acquired on a hairpin with a 15 nt loading site in the absence of roadblock. This overlap is quantitatively confirmed by the similar parameters that result from a bi-exponential fit to this re-merged distribution (*p*_1_ = 0.74 ± 0.10, *t*_1_ = 0.23 ± 0.09 *s* and *p*_2_ = 0.26 ± 0.10, *t*_2_ = 6.1 ± 2.9 *s*). In other words, despite the fact that the characteristic signature of the loose-lock binding mode on the hairpin hopping dynamics (Fig. 3A-C) is lost in the presence of the 5’-roadblock, as described in the preceding paragraph, its imprint on the kinetics of the start-up times is found to persist (Fig. 3F). This observation supports the hypothesis that a conformationally different (dsDNA-encircling) heptameric helicase competes with other oligomers for binding to the DNA hairpin at increased gp4 concentrations.

### Gp4 strongly interacts with the DNA fork directly downstream of the loading site

To further establish that the concentration dependence of the start-up times in the tight-lock configuration does not arise from the binding of multiple helicases, and also compare our experiment with previous single-molecule force spectroscopy studies of T7 gp4 helicase which typically employed loading sites of 400-600 nt ^13–15^, we studied the effect of a longer loading site. We select a hairpin with a 40 nt loading site (Fig. 4A), sufficiently large for (i) a single gp4 to bind even when the hairpin is fully closed and (iii) two gp4 to bind upon thermally-induced partial opening of the hairpin ^24^. As expected, this hairpin exhibits similar hopping dynamics at the base of its stem (red segments in Fig. 4B,C). We observe that the median time prior to helicase binding *T* is reduced to 28 ± 7 s for this hairpin (5 nM gp4, **Fig. S2J**), a (22 ± 6)-fold decrease compared to the hairpin with the 15 nt loading site. This reduction is not surprising, as adding an extra 25 nt to a 15 nt loading site effectively provides an additional 25 binding sites for gp4 to bind to.

In the presence of the longer loading site (Fig. 4A), we observe similar changes in hopping dynamics upon the addition of gp4, implying that helicase binding to the 40 nt loading site occurs at the fork. We continue to observe both binding modes (Fig. 4B, C, respectively), with the signature of the loose-lock configuration appearing at 5 nM on this construct, whereas this behavior required a gp4 concentration of 10 nM on the hairpin with the 15 nt loading site (Fig. 2-3). The fact that for a long loading site, binding still biases the hairpin hopping dynamics and we still observe both binding modes suggests a strong affinity of the initially loaded structures of both assembly pathways to the DNA fork. The presence of the tight-lock binding mode and, in particular, its similar start-up time kinetics (*t*_1_ = 0.6 ± 0.1 s, Fig. 4D) indicates that the stepwise assembly mechanism observed with the 15 nt loading site hairpin is maintained on a hairpin with this longer loading site and is unaffected by the potential proximity of additional gp4 oligomers upstream of the fork. In addition, this suggests that the oligomers that bind the ssDNA at the fork strongly enough to produce the tight-lock binding mode do so without needing to be ‘trapped’ on a short ssDNA loading site. The presence of the loose-lock binding mode provides additional support, in conjunction with the experiments on the hairpin containing a dsDNA roadblock, for the conclusion that this mode of binding does not necessarily require gp4 to overlap the dsDNA upstream of the loading site. We observe the distribution of start-up times τ for the loose-lock configuration (green data in Fig. 4E) to yield a shorter lifetime than in previous measurements, with the tight- and loose-lock configuration being governed by a single rate (0.6 ± 0.1 s and 0.7 ± 0.2 s, respectively) that is similar to the start-up rate for hexameric gp4 loaded onto the hairpin with the short loading site.

## Discussion

We have introduced a novel, label-free, force spectroscopy-based assay for probing DNA-protein interactions that makes use of a DNA hairpin construct that undergoes fast hopping dynamics on the first ~11 bp at the base of the hairpin stem at an applied force (~13.5 pN) just below the critical unwinding force (*F*_crit_ = ~ 16 pN). Using this assay, we detect protein binding from the changes brought about in the hopping dynamics at the base of the hairpin stem, and we detect subsequent enzymatic activity by monitoring protein translocation through the full stem of the hairpin.

We utilize this assay to characterize the loading mechanism of the T7 gp4 primase-helicase, a model hexameric replicative helicase of the superfamily IV. To do so, the DNA hairpin is preceded by a ssDNA loading site. At the tension applied, the extension of the ssDNA loading site is governed by entropic forces (as opposed to enthalpic forces), and hence the mean spacing between nucleotides along the direction of the pulling force remains well below the canonical interphosphate distance^30,31^. We observe that gp4 binding results in the hairpin becoming trapped in either a tight- (Fig. 2B) or loose-lock mode (Fig. 3B). These modes persist for a start-up time τ prior to gp4-driven unwinding of the complete DNA hairpin, with each mode being associated with a distinct mono-exponential distribution for τ (Fig. 3D, Fig. 3G, Fig. 4E). These observations are taken into account in the model displayed in Fig. 5, which we now proceed to describe in further detail.

**Fig. 5:**
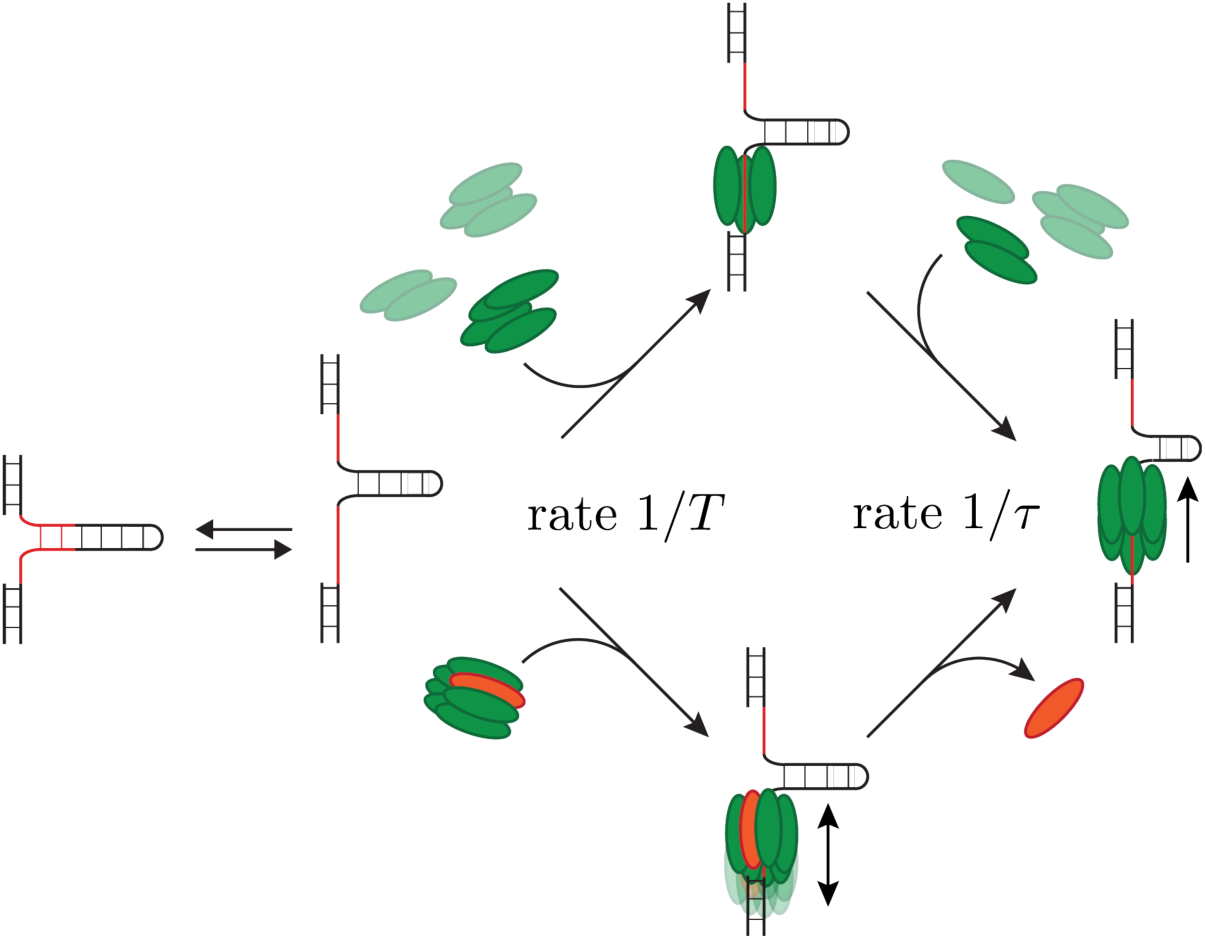
Model depicting the two distinct pathways through which T7 gp4 is loaded onto ssDNA. (Upper pathway) At low concentrations, oligomers smaller than a hexamer form the dominant conformation of gp4 in solution. These bind tightly to the ssDNA loading site (red) at a rate 1/*T* as shown; overall, we find that an unwinding-competent hexamer is assembled in a two-step process at a rate 1/ τ. (Lower pathway) At high concentrations, small oligomers coexist with a heptameric form of gp4 that binds loosely to the ssDNA loading site. Eventually, one of the monomers (orange) must disassemble at a rate 1/ τ for the unwinding-incompetent heptamer to mature into an unwinding-competent hexamer. In these two pathways, the start-up times τ need not be numerically identical.

For the tight-lock configuration, we are able to probe the concentration dependence of the start-up time and find it to decrease quadratically with increased gp4 concentration (Fig. 2E). Together with the mono-exponential distribution of start-up times, this indicates that gp4 assembly onto the ssDNA loading site at these concentrations occurs in at least two steps (upper pathway in Fig. 5), with the last step being a cooperative-binding step. It also suggests that even an incomplete gp4 ring can bind ssDNA sufficiently tightly to halt the hairpin hopping dynamics (data in Fig. 2C, upper pathway in Fig. 5). Such an iterative mode of assembly onto DNA, which is also reported for hexameric translocases (e.g. FtsK) and superfamily III hexameric helicases (e.g. SV40 large T antigen) ^9^, bypasses the need for spontaneous ring opening to achieve loading. We note that this type of assembly contradicts the results of previous studies on superfamily IV hexameric replicative helicases ^7,9,10^; however, it should be noted that these previous studies were performed at substantially higher gp4 concentrations (100 nM – 1 µM), at which gp4 is known to form large oligomers in solution ^24^.

To further establish that the concentration dependence in the start-up times does not arise from the binding of multiple helicases, we explicitly increase the probability of loading multiple helicases by studying a longer loading site. When the size of the ssDNA loading site is increased to 40 nt, we observe that the tight-lock binding mode is not eliminated (Fig. 4B), despite the loading site being sufficiently long to accommodate the presence of a single helicase while the base of the DNA hairpin stem is closed. The fact that the tight-lock persists indicates a strong preference for gp4 binding and/or assembly at the fork. Our results, supported by a study that investigated the affinity of gp4 for the DNA fork ^32^, suggest that the interactions between the gp4 helicase domain and the DNA fork are essential for gp4 binding and assembly onto ssDNA. We note that on such an extended loading site, a tight-lock configuration could hypothetically result from the adjacent binding of two hexameric gp4 molecules; however, given that the lifetime of the tight-lock state is 0.6 ± 0.1 s, whereas the median time to helicase binding is 28 ± 7 s (Fig. 4A), this seems unlikely. Furthermore, the lifetimes of the tight-lock binding modes on the 15 nt and 40 nt loading sites are in excellent agreement with one another (compare Fig. 2E and Fig. 4D, both acquired at 5 nM gp4). Taking these observations into account, we conclude that gp4 assembly, and not the loading of multiple hexameric gp4 molecules, forms the rate-limiting step during the start-up time that precedes unwinding.

A previously proposed mode for gp4 helicase binding to a ssDNA loading site is through a ring breaking mechanism of its heptameric form ^7^, followed by the loss of a monomer and the start of helicase activity. Such a heptamer-to-hexamer transition appears essential for tight coupling of gp4 onto the ssDNA followed by processive unwinding of the downstream dsDNA ^7^. The heptamer, with its relatively large inner channel of 2.5 nm diameter ^4^, could encircle the dsDNA directly upstream of the loading site, which could very well give rise to the reduced hairpin hopping rate of the loosely bound state (Fig. 3F). To probe the origin of the long loose-lock start-up times, we performed measurements with an additional short hairpin at the upstream boundary of the ssDNA-loading site, thereby hindering any backwards sliding. Introducing this roadblock, the probability to observe the loosely bound state indeed drops dramatically (Fig. 3F). Still, the distributions of start-up times remain unchanged (Fig. 3G), suggesting that the long start-up times in the absence of the roadblock are not due to the heptamer encircling the upstream dsDNA. Based on this, we suggest that the heptameric form does indeed load through a ring-opening mechanism^7,10^ (lower pathway in Fig. 5), giving rise to the loose-lock signature by encircling the dsDNA, and indicating that short ssDNA loading site is sufficient for heptamer loading. The continued presence of the loose-lock configuration on a DNA hairpin with a 40 nt loading site suggests that, as in the case of the tight-lock binding mode, the helicase remains in direct contact with the DNA fork. However, the strong reduction in the loose-lock start-up time (compare the green distributions in Fig. 4E and Fig. 3D) in this configuration suggests that the presence of a longer ssDNA loading site facilitates the conformational transition from heptamer to unwinding-competent hexamer. Within this context, we would predict that the start-up times associated with the loose-lock binding mode to be concentration-independent; however, the limited concentration range that is experimentally accessible prevents us from testing this directly.

We note that the existence of the tight- and loose-lock binding modes may also be interpreted differently, if one assumes that the gp4 hexamer can undergo a structural transition that allows its inner channel to modify its diameter. Such a conformational transition is observed for the *E.coli* replicative helicase dnaB ^33^. We deem such a mechanism unlikely though, as such a (stochastic) switching process should not be dependent on the concentration of gp4, in contrast to our observations. We thereby conclude that initiation by T7 gp4 proceeds via the two pathways described in Fig. 5. Given that T7 gp4 concentration *in vivo* is most likely far below the ~µM range of standard bulk experiments ^7,9,10^, we propose that the gradual helicase assembly pathway that we here detail for the first time is the most relevant during T7 genome replication *in vivo*.

## Acknowledgments

We thank Ivana Cvijovic and Tim Allertz for their contributions to initial datasets, Jelmer Cnossen for software interface development, and Theo van Laar for the synthesis of DNA hairpins. We thank Antoine van Oijen, Charles Richardson and Pauline van Nies for fruitful discussions. Funding for this work was provided by a TU Delft start-up grant to MD and by the European Research Council through a Consolidator Grant (DynGenome, N°312221) to NHD.

## Contributions

Designed the research: DD and NHD. Designed the single-molecule experiments: DD. Performed the experiments: ZY, DD and TJC. Analyzed the data: DD, ZY, and BAB. Interpreted the data and wrote the paper: DD, BAB, MD, and NHD.

